# The rapid divergence of the Antarctic crinoid species *Promachocrinus kerguelensis*

**DOI:** 10.1101/666248

**Authors:** Yacine Ben Chehida, Marc Eléaume, Cyril Gallut, Guillaume Achaz

## Abstract

Climatic oscillations in Antarctica caused a succession of expansion and reduction of the ice sheets covering the continental shelf of the Southern Ocean. For marine invertebrates, these successions are suspected to have driven allopatric speciation, endemism and the prevalence of cryptic species, leading to the so-called Antarctic ‘biodiversity pump’ hypothesis. Here we took advantage of the recent sampling effort influenced by the International Polar Year (2007-8) to test for the validity of this hypothesis for 1,797 samples of two recognized crinoid species: *Promachocrinus kerguelensis* and *Florometra mawsoni*. Species delimitation analysis identified seven phylogroups. As previously suggested, *Promachocrinus kerguelensis* forms a complex of six cryptic species. Conversely, despite the morphological differences, our results show that *Florometra mawsoni* is a lineage nested within *Promachocrinus kerguelensis*. It suggests that *Florometra mawsoni* and *Promachocrinus kerguelensis* belong to the same complex of species. Furthermore, this study indicates that over time and space the different sectors of the Southern Ocean show a remarkable rapid turn-over in term of phylogroups composition and also of genetic variants within phylogroups. We argue that strong “apparent” genetic drift causes this rapid genetic turn-over. Finally, we dated the last common ancestor of all phylogroups at less than 1,000 years, raising doubts on the relevance of the Antarctic “biodiversity pump” for this complex of species. This work is a first step towards a better understanding of how life is diversifying in the Southern Ocean.

## Introduction

During the last decade following the last International Polar Year (2007-8) a huge effort has been made to better understand Antarctic marine ecosystems and one of the major outcomes has been the creation of a reference barcode database (BOLD) that reconciles morphologically recognized species with a unique COI fragment. This opened to a new era of discoveries that allowed reassess some of the main known characteristics of marine Antarctic ecosystems, including eurybathy, excess in brooding species, high rate of endemism, rapid diversification and cryptic speciation as a consequence of the biodiversity pump (Clarke & Crame, 1989). Recent biodiversity assessments using molecular tools have revealed an increasing number of cryptic species in teleost fishes as well as Echinodermata, Mollusca, Arthropoda, or Annelida (see (Cornils & Held, 2014; Griffiths, 2010; Rogers, 2007) for comprehensive reviews). Cryptic species have been shown to be homogeneously distributed among taxa and biogeographic regions (Pfenninger & Schwenk, 2007) and Antarctica may well be no exception.

The Southern Ocean is known to have undergone several glaciation events (Clarke & Crame, 1989; Clarke, Crame, Stromberg, & Barker, 1992; Thatje, Hillenbrand, Mackensen, & Larter, 2008). Massive ice sheet advance and retreat seem to have bulldozed the entire continental shelf several times during the last 25 Mya. These events are thought to be driven by Milankovitch cycles. These cycles may be among the strongest evolutionary forces that have shaped Antarctic terrestrial and marine biodiversity (Clarke et al., 1992; Thatje et al., 2008; Thatje, Hillenbrand, & Larter, 2005). Thatje *et al.* (2005; 2008) hypothesized that vicariant speciation could have occurred during the glacial periods on the Antarctic continental shelf, within multiple ice-free refugia like polynya. As lineages evolved independently during the glacial period, they accumulated genetic differences that lead to reproductive isolation and probably to the formation of cryptic species. During interglacial periods, barriers to gene flow may have been removed allowing for secondary contact between the vicarious lineages (Heimeier, Lavery, & Sewell, 2010; Thatje et al., 2005; 2008; Thornhill, Mahon, Norenburg, & Halanych, 2008). An alternative hypothesis is based on the idea that Antarctic continental shelf may be understood as a species flock generator (Eastman & McCune, 2000; Lecointre et al., 2013).

Here, we have analyzed COI sequences of 1,797 individuals of two crinoid species: *Promachocrinus kerguelensis* (Carpenter, 1888) and *Florometra mawsoni* (AH Clark, 1937), endemic to the Southern Ocean. These species represent the most abundant crinoid species in the Southern Ocean (Eléaume, Hemery, Roux, & Améziane, 2014; Speel & Dearborn, 1983). They are morphologically distinct (but see (Eléaume, 2006) for counter arguments) and genetically close (Hemery, Améziane, & Eléaume, 2013a). These species are thought to have a reproduction cycle that involves external fertilization that could result in a large dispersal potential. *P. kerguelensis* produces positively buoyant ovocytes that, after fertilization, may remain in the plankton for weeks or even months (McClintock & Pearse, 1987). Some adults of *P. kerguelensis* have also been observed swimming (Eléaume, unpublished observations), and some adults of *F. mawsoni*, though never observed swimming in situ, have shown this ability in tanks (Eléaume, unpublished observations).

During the last decade, numerous specimens of crinoids have been collected from all around Antarctica and sub-Antarctic islands. Over 3,000 specimens attributed to 45 species have been sampled (Eléaume et al., 2014). Species such as *Promachocrinus kerguelensis* and *Florometra mawsoni* are represented by a large number of well distributed specimens. Knowlton (1993; 2000) predicted for Southern Ocean organisms that an increase in sampling effort and the application of genetic tools, would reveal cryptic species. The analysis of *P. kerguelensis* COI barcode fragment suggested that this species may constitute such an example of a complex of unrecognized species. Wilson *et al.* (2007) identified six lineages in the Atlantic sector whereas Hemery *et al.* (2012) extending the analysis of Wilson to the entire Southern Ocean, identified seven lineages. As no obvious morphological character have been shown to distinguish between these lineages (Eléaume, 2006), they may represent true cryptic species. However, Hemery *et al.* (2012) using nuclear markers have shown that the six identified COI lineages may only represent three distinct entities. All of the six or seven lineages are circumpolar in distribution, sympatric and eurybathic but show various levels of connectivity that depend on the lineage and the geographical area.

Here we analyze the pooled COI datasets of *P. kerguelensis* and *F. mawsoni* of Wilson *et al.* (2007) and Hemery *et al.* (2012) using different approaches to separate different sources of genetic variation, i.e. differences due to diversification (clade), time (year collected), space (geographical origin of samples). We first analyze the phylogenetic relationships within *P. kerguelensis* sensu largo using different species delineation methods. We then measure the genetic turn-over, within each phylogroup sampled at a given location. Using a population genetics coalescent framework, we estimated an extraordinary small effective population size that corresponds to a mutation rate that is larger than previously reported before. We then discuss our results in the light of the climate history of the Southern Ocean.

## Materials and methods

### Sequences

We included 13 sequences of *P. kerguelensis* from Wilson *et al.* (2007), for which we had precise information about the place and the date of sampling. We have also included 1303 sequences of *P. kerguelensis* from Hemery *et al.* (2012) together with 479 sequences of *F. mawsoni* (Hemery *et al.*, in preparation). A total of 1,797 COI sequences were used for this study. All specimens were collected between 1996 and 2010 in seven geographical regions that were described in Hemery *et al.* (Hemery et al., 2012): Kerguelen Plateau (KP), Davis Sea (DS), Terre Adélie (TA), Ross Sea (RS), Amundsen Sea (AS), West Antarctic Peninsula (WAP), East Weddell Sea (EWS). We however further split the Scotia Arc into Scotia Arc East (SAE) and Scotia Arc West (SAW) as there can be as much as 2,000 km between them. The counts of all sequences of all locations are reported in Table S1.

### DNA extraction, PCR and sequencing

For this analysis, no additional sequences have been produced. For DNA extraction and PCR procedures see (Hemery et al., 2012; Wilson et al., 2007) and 2013. A total of 1797 554-bp sequences of the barcode region of cytochrome c oxidase subunit I (COI) were amplified using the Folmer *et al.* (1994) primers and other specific primers described in Hemery *et al*. (2012). All COI sequences from Hemery *et al.* (2012) have been made easily available through a data paper article (Hemery et al., 2013a).

### Species delimitation

We used three independent species delimitation methods: Generalized Mixed Yule Coalescent (GYMC, (Pons et al., 2006)), Automatic Barcode Gap Discovery (ABGD, (Puillandre, Lambert, Brouillet, & Achaz, 2011)) and Poisson Tree Process (PTP, (Zhang, Kapli, Pavlidis, & Stamatakis, 2013)) to estimate the number of phylogroups in our sample. For all these methods, we used all the 199 unique haplotypes of the dataset to have legible results and a faster computation time. The phylogenetic tree used for PTP was constructed by Neighbor Joining method the BioNJ implementation (Gascuel, 1997) in *Seaview* software (Gouy, Guindon, & Gascuel, 2010) version 3.2. PTP analysis was conducted online (http://species.h-its.org/ptp/). The ultrametric tree needed for GYMC method was constructed using *BEAST* software (Bayesian Evolutionary Analysis Sampling Trees, (Drummond, Suchard, Xie, & Rambaut, 2012) version 1.7). GYMC analysis was conducted online (http://species.h-its.org/gmyc/). ABGD analysis was conducted online (http://wwwabi.snv.jussieu.fr/public/abgd/abgdweb.html).

### Assessing the temporal structure within phylogroups

We characterized the temporal structures of our samples in pairwise comparisons using both fixation index *F_ST_* as well as a non-parametric structure test based on *K*\**s* by Hudson *et al.* (1992). All p-values were estimated using permutations as described in Hudson *et al.* (1992). A web-interface for the latter can be found at http://wwwabi.snv.jussieu.fr/public/mpweb/.

### Estimation of the mutation rate and the population effective size

As described by Fu (2001) one can use temporal data to estimate the mutation rate (*μ*). In summary, the method assumes that the population is sampled in at least two time points (say *T*_*1*_ and then later *T*_*2*_) with multiple sequences in *T*_*1*_. The expected average pairwise differences between sequences taken from the two time points (*K*_*12*_) equals the average pairwise differences of sequences within the past time point (*K*_*11*_) plus the difference accumulated since then. Mathematically, it is expressed as *E[K*_*12*_] = *E[K*_*11*_] + *(T*_*2*_*−T*_*1*_*)μ*. Because the time difference between the samples (*T*_*2*_*−T*_*1*_) is known, the mutation rate can be estimated from the observed pairwise differences: *μ* = *(K*_*12*_*−K*_*11*_)/*(T*_*2*_*−T*_*1*_). This idea can be expanded when more data are available in other time points (Fu, 2001).

Once μ is known, the average pairwise difference can be used to estimate the effective population size (*Ne*). Under the standard neutral model, the pairwise difference within groups of the same sample equals *2Neμ*, when *μ* is expressed as a rate by generations. Here, we assumed one generation every three year to estimate *Ne* (Clark, 1921).

### Estimation of the time of divergence between phylogroups

For visualization purposes, an ultrametric tree was constructed with the unique sequences by UPMGA (Michener & Sokal, 1957) as implemented in the *Phylip* package (Felsenstein, 1989) version 3.6. Using the estimated mutation rate, we derived a time of divergence within and between phylogroups from the average sequence differences.

### Polymorphisms analysis

We used *Dnasp* software version 5.10 (Librado & Rozas, 2009) and the program used to study genetic structure to study the SFS (Site Frequency Spectrum) within phylogroups as well as two of its summary statistics: nucleotide diversity (π) and number of polymorphic sites (S). We then tested neutrality using Tajima’s D (Tajima, 1989), Fu and Li’s F* and D* (Fu & Li, 1993), Achaz’s Y* (Achaz, 2008), that is immune to sequencing errors, and the Strobeck (Strobeck, 1987) haplotype test.

### Assessment of migration effect on Ne estimation through simulations

To assess the potential effect of migrants from a distant population on the estimation of *μ* and *Ne*, we used coalescent simulations where two samples of 20 sequences each are taken 3N generations apart in a focal population of size *N*. We assume a large unsampled ghost population of size *10N* that exchange migrants with the focal one at rate *M=4Nm*, where *m* is the migration rate per generation. For values of *M* ranging from 0.001 (very low) to 1000 (very large), we applied the Fu (Fu, 2001) method to estimate *Ne*, with 10^3^ replicates for each *M* value. The program is based on a C++ library developed by G. Achaz that is available upon request.

## Results

### P. kerguelensis and F. mawsoni are composed of seven phylogroups

We first reinvestigated the species delimitation of the sample using three independent approaches (GMYC, ABGD and PTP) that cluster the unique sequences by phylogroup. We adopt in this manuscript a nominalist definition of species. However, *F. mawsoni*, a nominal species, is nested within *P. kerguelensis*, another nominal species, lineages. To avoid confusion, we have decided to use the term “phylogroup” in the rest of the manuscript to designate any well-supported cluster of sequences derived from species delineation approaches.

Two of the three methods estimate a similar number of phylogroups: PTP estimates that there are nine phylogroups in the sample whereas ABGD reports 5-7 phylogroups, depending on the chosen prior value. Looking at the proposed partitions, we have chosen to consider seven different phylogroups for this study that are reported as phylogroups A-G in Fig. 1. Interestingly, these phylogroups correspond to the A-F groups proposed by Wilson *et al.* (2007), plus a ‘new’ group (G) that corresponds to the *F. mawsoni* species. The two extra sub-splits suggested by the PTP method are indicated as C1/C2 and E1/E2 on Fig. 1. Note that PTP only seeks monophyletic groups and thus must split the C group into the C1/C2 subgroups.

**Fig. 1:**
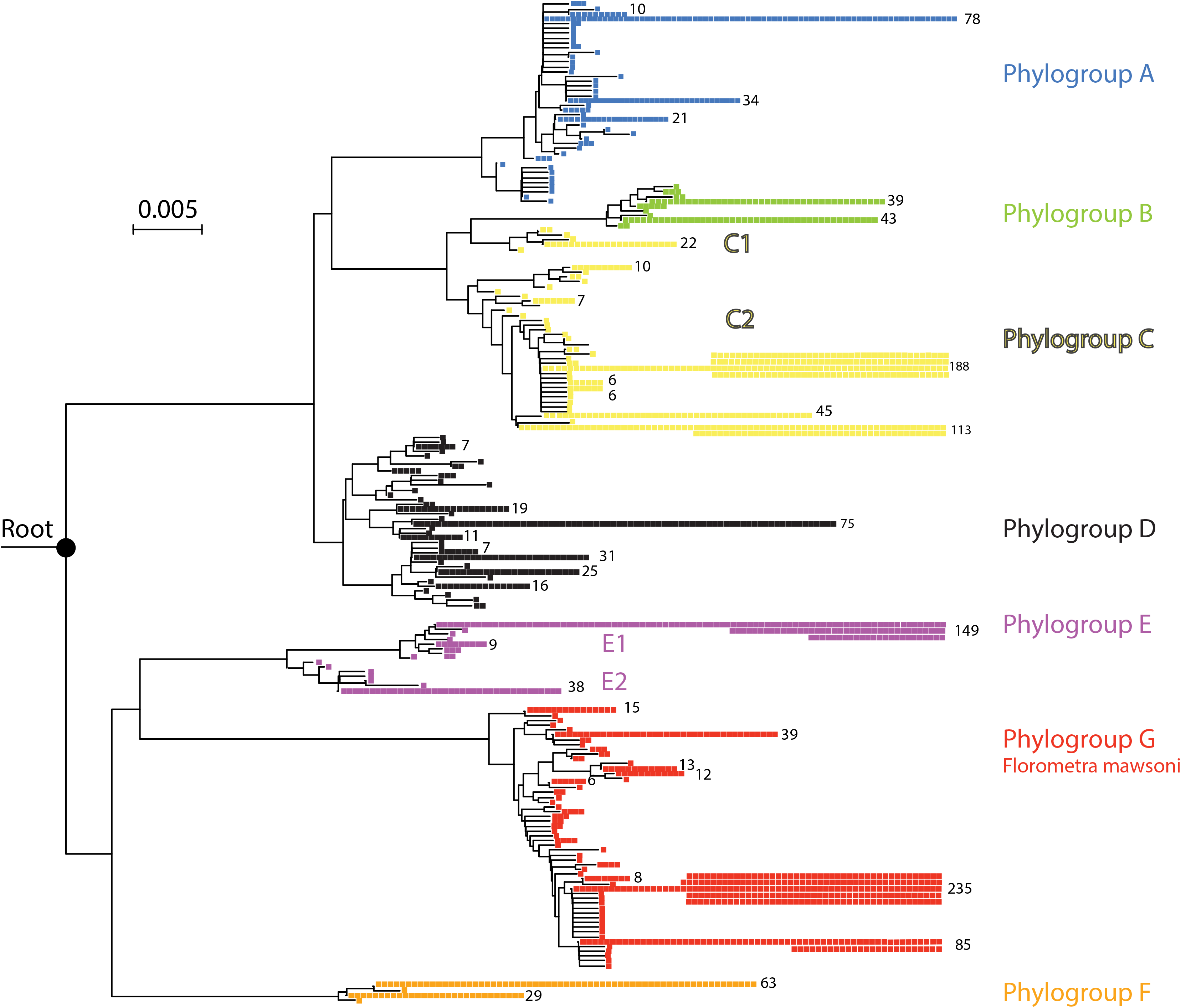
Phylogenetic representation of the 1797 COI sequences from *P. kerguelensis* and *F. mawsoni*. The tree was inferred using BIONJ on the unique sequences. Each square represents one sequence. The phylogroups are the ones used in the analysis (mostly ABGD groups) and both supplementary splits proposed by PTP are indicated as C1/C2 and E1/E2. The colors represent the different phylogroups considered in this study. The boxes and the numbers refer to the number of individuals carrying the haplotype.

In contrast with the two previous methods, depending on the sequence evolution model we used and the *BEAST* parameters, GMYC reports between 3 and 51 phylogroups with non-overlapping grouping between individuals. As GMYC is very sensitive in our case to the chosen parameters, it is here unreliable and we chose to not consider the GMYC partitions.

Genetic distance between phylogroups are similar to the distance of each phylogroup to the G group. However, G group corresponds to a nominal species (*F. mawsoni)* which suggests that the A-G phylogroups could correspond to seven different yet undescribed species.

### A very rapid local turn-over

#### Relative abundance of phylogroups

Two of the analyzed locations (Ross Sea and East Weddell Sea) were densely sampled (at least 50 sequences) at two (or more) different time points. We therefore took advantage of these time series to evaluate the turnover of the resident crinoids from year to year. The analysis of the relative abundance of the various phylogroups shows that for both locations the genetic composition is highly unstable through time (Fig. 2a and 2c). Phylogroup composition in EWS is stable from 1996 to 2004, and changes drastically in 2005 when phylogroup C has not been sampled. This pattern leads to a highly significant difference between all years (Fisher exact test, P = 4.7 10^−10^). Similarly, the composition of the RS location is also highly significantly different between 2004 and 2008 (Fisher exact test, P = 3.2 10^−11^); in this case, phylogroup A is replaced by phylogroups B and E.

**Fig. 2:**
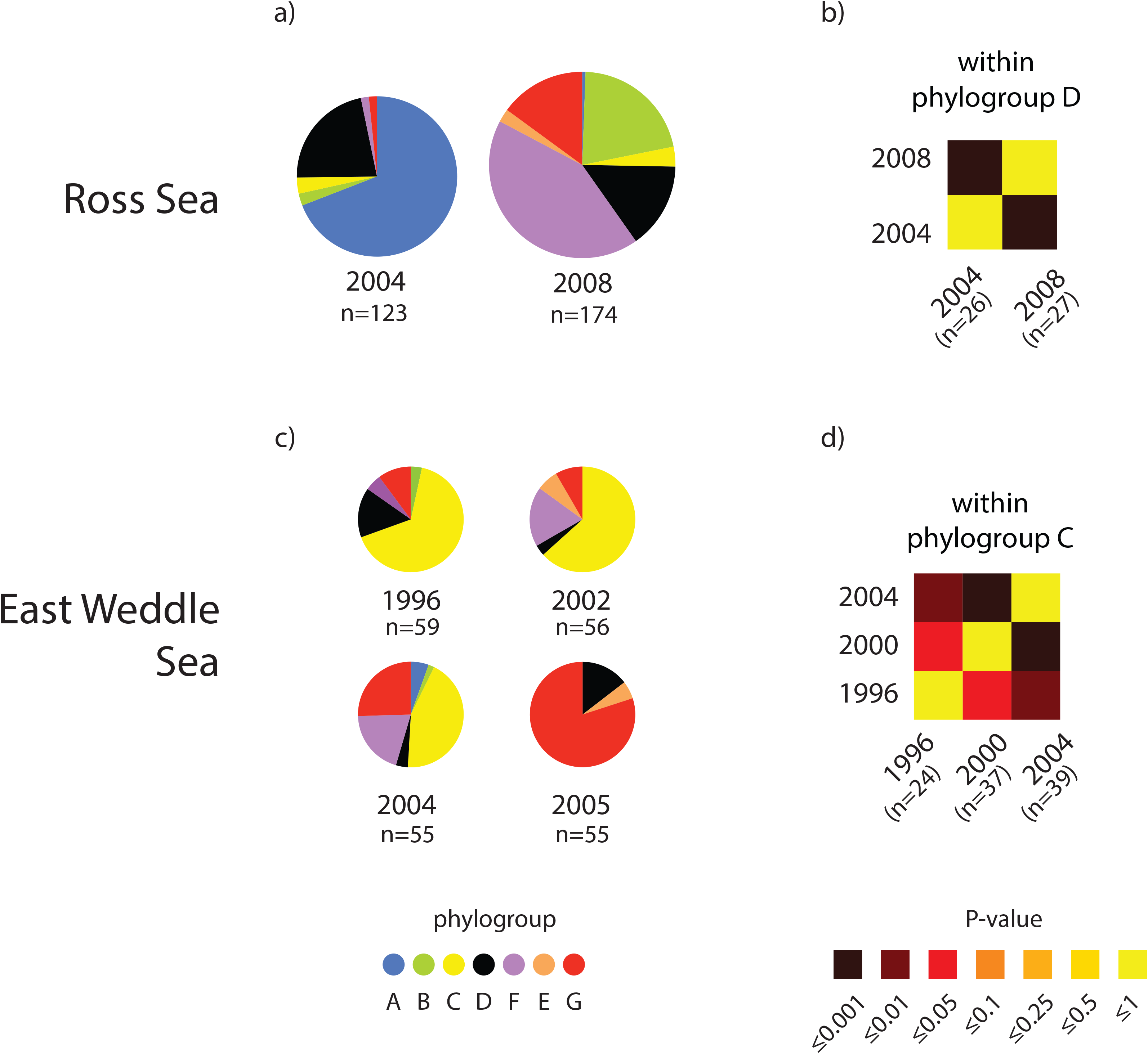
Genetic turn over at the RS (a and b) and EWS (c and d) locations. The pie charts (a and c) represent the number of phylogroups at different years in the locations, with the same colors code as in Figure 1. The heatmaps (b and d) represents P-values of the Hudson et *al.* (1992) tests for genetic structure within a phylogroup of each location.

Other locations display a single sampling event (AS, DS, TA) or reduced number of sequences over several sampling events (e.g. SAW: 2 sequences in 2000, 11 in 2002, 7 in 2006 and 68 in 2009), and cannot be used to robustly assess the stability of phylogroup relative abundance.

#### Turn-over within the phylogroups

We then measured the replacement of genetic variants within the phylogroups. To do so, we considered samples with at least 10 sequences of the same phylogroup in the same location at two different time points. The pairwise comparisons for such groups are all reported in Table 1 and we provide two graphical representations for phylogroup D in the RS location (Fig. 2b) and phylogroup C in the EWS location (Fig. 2d). In two cases (phylogroup C in EWS and phylogroup D in RS), we found a significant difference in the genetic composition within each phylogroups even when they are separated by only few years, again suggesting an unexpectedly high turn-over of the genetic pool of these crinoids. Interestingly, the temporal differentiation of phylogroup E within EWS is already *F*_*ST*_=4% in four years, that is not significant because of the small sample size (10 and 11 respectively). Only the two samples of phylogroup G in EWS, that are separated by a single year, show no sign of temporal differentiation (*F*_*ST*_=1%).

**Table 1:**
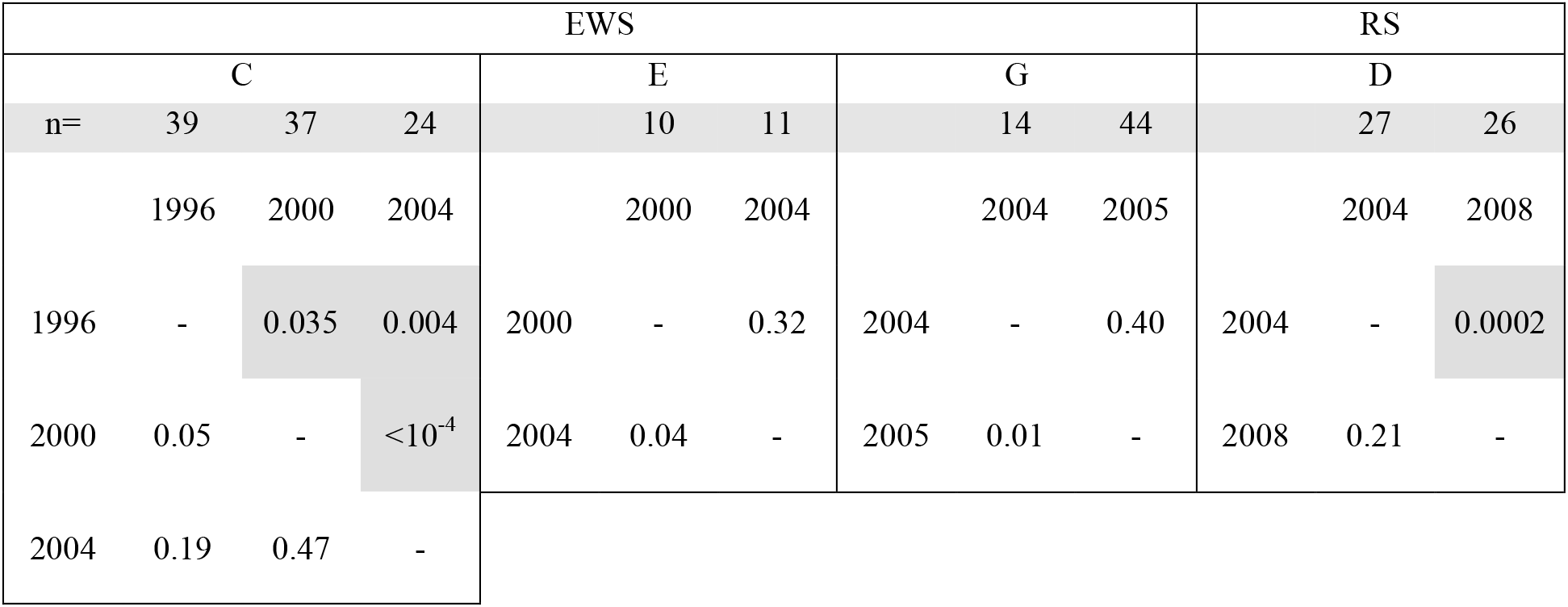
Pairwise F-st and the probability from the Hudson et al. (1992) tests for all phylogroups with 2 samples of 10+ sequences within the same location. Below diagonal: Fst. Above diagonal: probability of pamixia using the Hudson test.

### Estimation of the mutation rate and the effective population size

Using the robust yet elegant approach devised by Fu (2001), we estimated the yearly mutation rates (μ) from the same sample comparisons (i.e. phylogroups with two or more samples of 10 or more sequences within the same location). Furthermore, we also estimated the corresponding effective population size (N_e_) assuming a generation time of three years. Results (Table 2) shows that all estimated mutation rates are 10^−5^-10^−4^ /bp/year, with a mean of 8.10^−5^ /bp/year. Results also show that the effective sizes are very low, on the order of 2-7 (Table 2). The very low effective size corresponds to the unexpected high turn-over of the genetic pool that we observe within the phylogroup at the same location.

**Table 2:** Estimation of mutation rate and effective population size. Summary table of effective population sizes (*Ne* in numbers of individuals) and mutation rates (μ per base pair per year) for each phylogroup estimated by the Fu (2001) method. We also report the years and the number sequences used for each estimation.

| Location | EWS |  |  |  |  |  |  |  | RS |  |
| --- | --- | --- | --- | --- | --- | --- | --- | --- | --- | --- |
| Phylogroup | C |  |  | E |  | G |  | D |  |  |
| n= | 39 | 37 | 24 | 10 | 11 | 14 | 44 | 27 | 26 |  |
| Dates | 1996 | 2000 | 2004 | 2000 | 2004 | 2004 | 2005 | 2004 | 2008 |  |
| $\mu$ | 1.9 10<sup>-5</sup> | | | 2.7 10<sup>-5</sup> | | 3.5 10<sup>-4</sup> | | 4.2 10<sup>-4</sup> | | |
| Ne | 4.0 |  |  | 7.0 |  | 1.7 |  | 2.6 |  |  |

### Polymorphism analysis

For samples of 10 sequences or more (with the same year and same location), we have analyzed the characteristics of segregating polymorphisms using four neutrality tests: Tajima’s D (Tajima, 1989), Fu and Li’s F* and D* (Fu & Li, 1993), Achaz’s Y* (Achaz, 2008) and Strobeck P-values (Strobeck, 1987). Results (Fig. 3) show almost no deviation from the standard neutral model, except for the Fu and Li’s F* and D*. This statistics points to a clear excess of singletons. However, singletons likely contain sequencing errors that strongly skew these statistics (Achaz, 2008). The Y statistics that ignores singletons exhibits a distribution that is very close to the one expected under the standard neutral model. We therefore conclude that the observed polymorphisms suggest that basic assumptions of the standard neutral model (i.e. constant population size, no migration, no structure and no selection) cannot be rejected for these crinoid phylogroups.

**Fig. 3:**
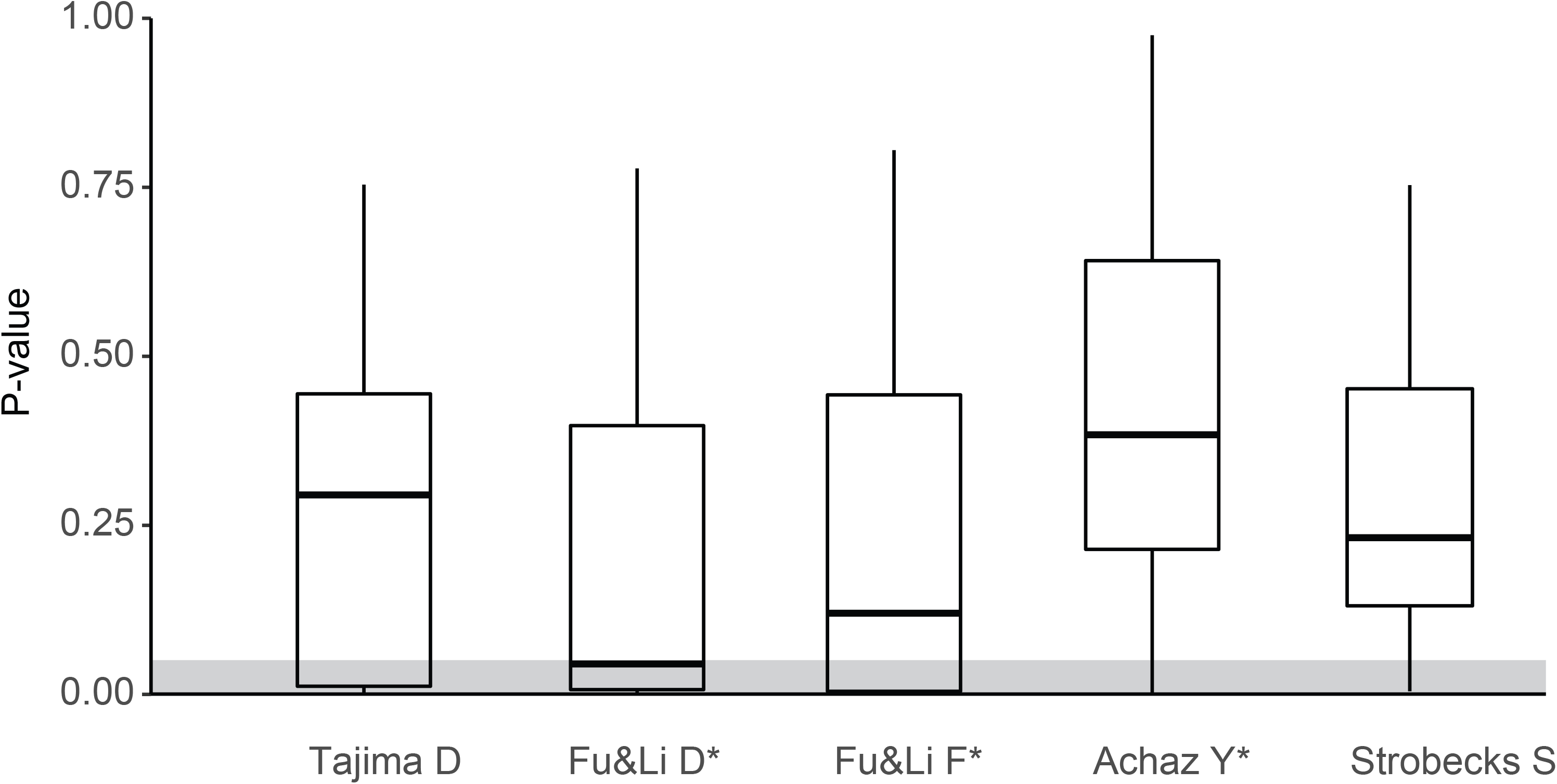
Distribution of p-values for different neutrality tests on the 1,624 sequences tested for group with at least 10 sequences. From left to right: Tajima D (1983), Fu and Li D* (1993), Fu and Li F* (1993), Achaz Y* (2008) and Strobeck S (1987).

### Assessing the effect of migration using simulations

To test whether migration could accelerate the genetic turn-over of a species on a given location, we studied a simple model where the sampled location exchanges migrants with a large pool of panmictic individuals. One expects that a constant arrival of migrants from this large pool could interfere with the estimation of effective size. We thus simulated a model where the sampled population has a size *N* and the pool 10*N*. We studied the impact of the migration rate on the estimation of Ne using the same method than above (2001). Results (Fig. 4) show that at low migration rate, the estimator is unaffected whereas it is inflated up to 11*N* at high migration rate. This shows that the existence of a large pool from where migrants can come into the sampled population can only inflate the estimation of *Ne* and therefore cannot explain the low *Ne* we have estimated.

**Fig. 4:**
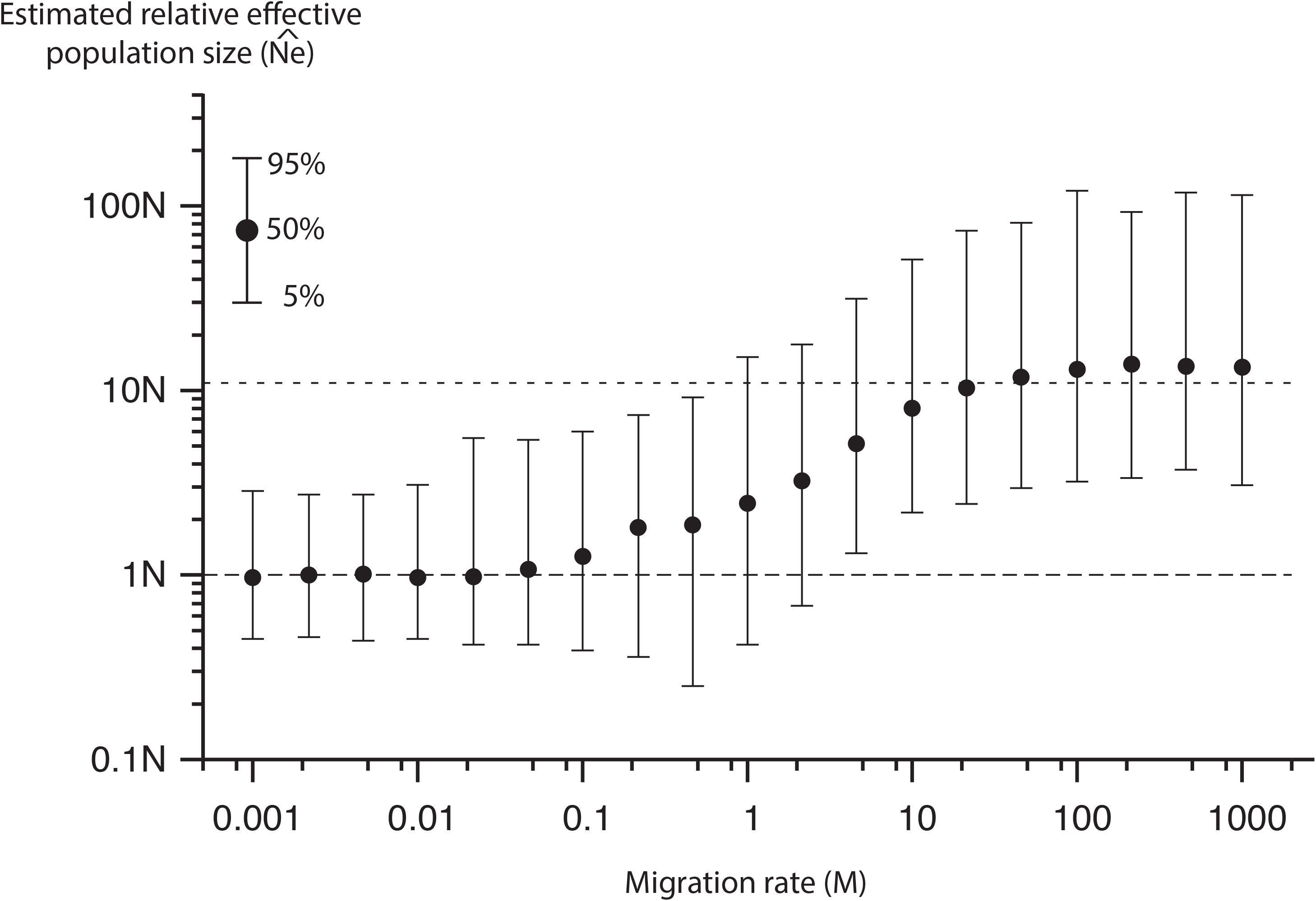
Simulation of migration effect on Ne estimation.

### Phylogroups divergence

We then computed an estimate for the times of split between the seven phylogroups using our population genetics estimation of the mutation rate (8.10^−5^ mutations /bp /year). Our estimates are reported in an ultrametric tree (Fig. 5) where the oldest split between the phylogroups is estimated around 400 years. This very short time scale is in sharp contrast with the millions of years of previous estimations based on other estimated mutation rates (3.10^−8^ mutations /bp /year) that were based on echinoderms thought to be separated by the formation of the panama isthmus (Lessios, Kessing, Robertson, & Paulay, 1999).

**Fig. 5:**
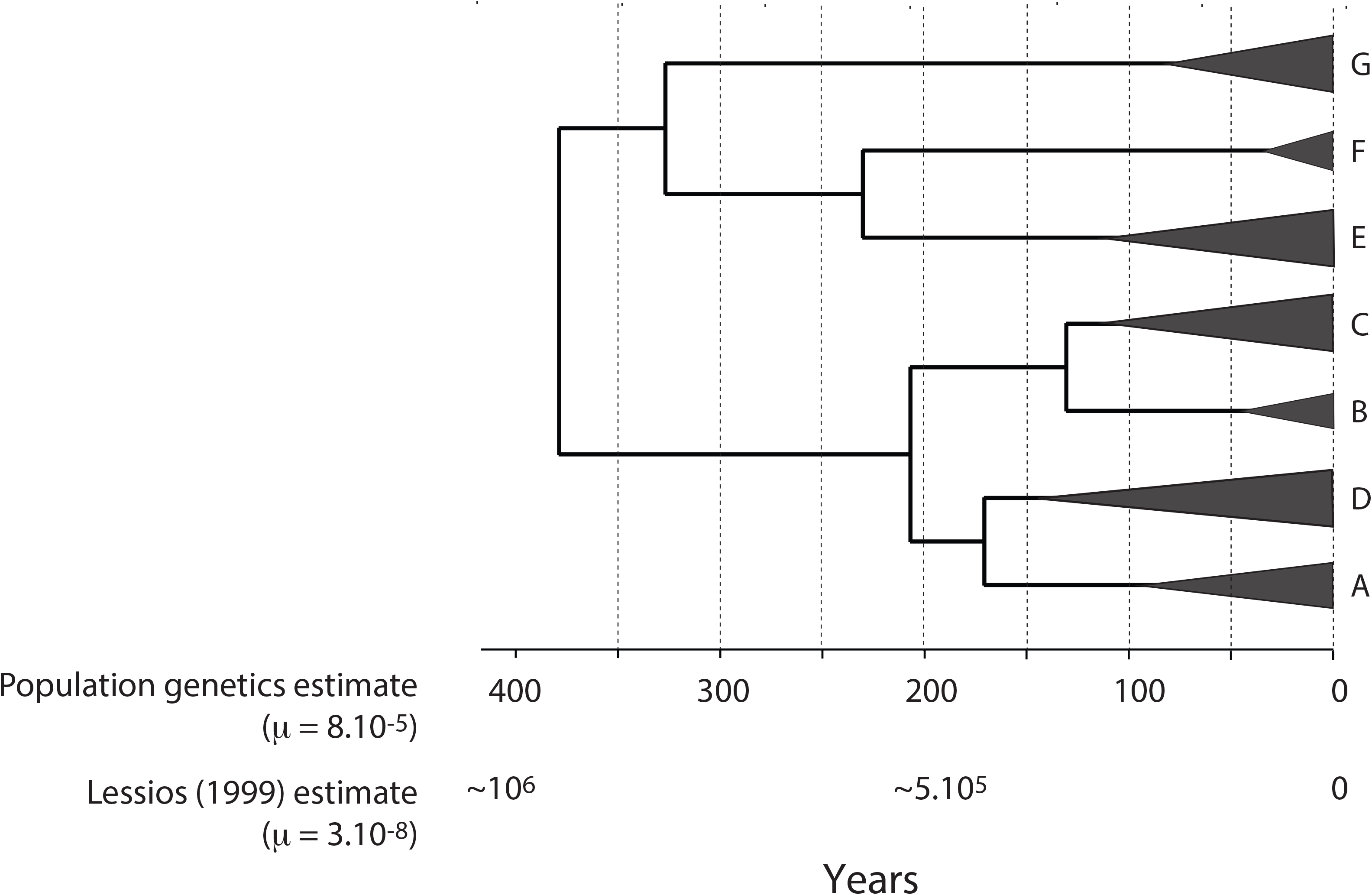
Divergence time of the different phylogroup of the *P. kerguelensis* complex. The depth of the triangles represents the maximum diversity within each phylogroup. The time is indicated in years before present. The timescale on the top is assuming a mutation rate of 8.10^−5^ mutation/bp/year. The timescale on the bottom is assuming a mutation rate of 8.10^−3^ mutation/bp/year.

## Discussion

### Speciation in Antarctica: the classical view

Due to its extreme environmental conditions, it has been hypothesized that climate driven evolution in Southern Ocean has been more important than in other ecosystems where ecological interactions play a larger role (Rogers, 2012). The diversity and the distribution of many marine taxa in Antarctica has been strongly shaped by climate variation and continental drift leading to dispersal, vicariance and extinction since the break-up of Gondwana (Williams, Reid, & Littlewood, 2003). Indeed, the global climate change at the end of the Eocene related to the continental drift is characteristic of the transition from a temperate climate to the current polar climate in Antarctica (Clarke et al., 1992; Clarke & Crame, 1989). The physical separation of Antarctica, South America and Australia resulted in the cooling of the Southern Ocean, which in turn initiated the physical and climate isolation of this region of the world (Tripati, Backman, Elderfield, & Ferretti, 2005). This isolation was followed by an extensive continental glaciations and the initiation of the Antarctic Circumpolar Current in the Miocene (Clarke et al., 1992; Clarke & Crame, 1989; Thatje et al., 2008). These two processes exacerbated the environmental and biological isolation of the Southern Ocean and are classically recognized as the main explanation to the high level of endemism observed in Antarctica (Clarke & Crame, 1989; Griffiths, Barnes, & Linse, 2009).

The Quaternary was marked by the oscillation of glaciation and interglacial periods, which strongly affected the ice cover present in Antarctica (Davies, Hambrey, Smellie, Carrivick, & Glasser, 2012). These cycles led to severe temperature oscillations and to the succession of large extensions of the ice sheets on the continental shelf followed by retreats of the ice sheets. They have been one of the main drivers of the diversification of species in Antarctica. Indeed, the isolation of populations in ice free refugia during the glacial era, such as polynya or in the deep sea, has been suggested as a mechanism of fragmentation of populations leading to allopatric speciation in the Southern Ocean. This process is called the Antarctic ‘biodiversity pump’ (Clarke et al., 1992; Clarke & Crame, 1989). The Antarctic ‘biodiversity pump’ has been proposed to be the main mechanism driving speciation and diversification in Antarctica.

Our knowledge on evolutionary history of Antarctic fauna was classically derived from studies of the systematics and distribution of marine animals relying heavily on the morphology of present and past (fossil) organisms. Such approaches are limited for two main reasons. First, species morphology may not reflect the true evolutionary relationships between taxa (DeBiasse & Hellberg, 2015). Second, several periods of time lack a reliable fossil record. Today, the emergence of molecular approaches provides a powerful tool to circumvent these limitations. To this regard, the increasing number of phylogeographic studies allow to get a better understanding of the effects of past changes on the current diversity and spatial distribution of organisms (Allcock & Strugnell, 2012). They also allow to accurately assess the distribution of genetic diversity within and among species. Such information is extremely valuable to evaluate and mitigate the impacts of human-induced activities on the biodiversity of the Southern Ocean (Chown et al., 2015). This study take place in this context and by expanding the results of two previous studies (Hemery et al., 2012; Wilson et al., 2007), we aimed to determine the drivers of the diversity of two crinoid taxa in the Southern Ocean, *P. kerguelensis* and *F. mawsoni*.

### The concept of species: is Promachocrinus kerguelensis a complex of multiple cryptic species?

The analysis of 1797 sequences using recent species delimitation approaches allowed to identify seven operational taxonomic units. We confirmed the presence of six phylogroups (A-F) identified in two previous study (Hemery et al., 2012; Wilson et al., 2007), but only the PTP method supports the further split between E1 and E2 suggested by Hemery *et al.* (2012). Furthermore, despite their strong morphological differences and monophyly (Fig. 1), our results strongly suggest that *F. mawsoni* (phylogroup G) belongs to the *P. kerguelensis* complex. Indeed, the reciprocal monophyly of G and F and the proximity of E to G/F compare to A-D confirm the affiliation of G as a member of *P. kerguelensis* complex (Fig. 1). Therefore, we corroborated Hemery *et al.* (2012) results showing that *P. kerguelensis* consists in a complex of several cryptic species. We also confirmed Eléaume (2006) conclusion that despite striking exterior morphological differences phylogroup G is also a member of this complex of species, and Hemery et al. (2013b) results based on a multi-markers phylogenetic reconstruction.

Morphological similarities between cryptic species often reflect a recent speciation event (Bickford et al., 2007). In the marine realm, speciation is less associated to morphology than to other phenotypic aspects such as chemical recognition systems (Palumbi, 1994). Knowlton (1993) argued that marine habitats are likely to be filled with cryptic species although rarely recognized due to limited access to marine habitats. Molecular approaches can be particularly powerful in detecting cryptic species on the basis of their molecular divergence, and recent molecular studies helped reveal the prevalence of cryptic species (See for example (Brasier et al., 2016; Grant, Griffiths, Steinke, Wadley, & Linse, 2010)). This is particularly the case in the Southern Ocean where numerous molecular studies have suggested the presence of cryptic species (Baird, Miller, & Stark, 2011; Brasier et al., 2016; Grant et al., 2010; Wilson, Schrödl, & Halanych, 2009) with areas such as the Scotia Arc representing potential hotspots of cryptic diversity (Linse, Cope, Lörz, & Sands, 2007). The prevalence of invertebrate cryptic species in the Southern Ocean is probably due to the cyclical variation of ice sheets extent in Antarctica during repeated glacial and interglacial periods which could have fostered the separation of population, promoting genetic divergence and allopatric cryptic speciation (Clarke & Crame, 1989). During glacial periods, ice sheets extension is thought to have forced marine species inhabiting the continental shelf to take refuge in the deep sea or in shelf refugia, such as areas where permanent polynyas (Thatje et al., 2008) occurred. During glacial maxima the decrease of gene flow between populations would have promoted reproductive isolation and increased genetic variation between populations. Under glacial maximum extreme environmental conditions, a potential increase of the diverging selective pressures on behavioral and physiological characters rather than morphology (which could be under strong stabilizing selection), could lead to a reduction of the morphological changes usually associated with speciation (Fišer, Robinson, & Malard, 2018). In such a scenario, high levels of divergence are expected on neutrally evolving genes as observed in this study. This process might explain the high number of cryptic species reported in the Southern Ocean.

The concept of species is controversial in biology as there is no unique definition that can be applied to all species under all circumstances (Mayden, 1997). Here, the application of species delimitation methods (based on the genetic concept of species) using molecular tools identified six highly divergent lineages indiscernible on the basis of their morphology and a last morphologically divergent lineage. Therefore, this study show how the use of molecular data can provide new insights into the nature of the genetic boundaries between species (Pante et al., 2015). However, in the light of these findings we can wonder what do the lineages highlighted in this study represent? Are they genuine different species? Here the use of different criteria to delineate species based on reproductive isolation (biological concepts of species) or on morphology (morphological concept of species) will probably lead to the identification of a different number of species (Fišer et al., 2018). This is a good illustration of the “species problem” and stress out the fact that the notion of species itself is an abstract concept that should be questioned (De Queiroz, 2007). It has also important consequences in term of the protection of the biodiversity because disparate groupings of species will in turn result in different economic implications and conservation decisions (Frankham et al., 2012).

### A remarkably fast turn over and low effective population sizes

This study indicates that over time the different sectors of the Southern Ocean show a remarkable turn over in term of phylogroup composition. For example, in less than a decade, the Ross and East Weddell Sea phylogroups composition changed substantially (Fig 2a and 2c). A, B and F may or may not be sampled from one year to the next, whereas C, D and G are always present but their proportions change drastically (Fig 2a and 2c). This observation suggests that within the same geographical region, the real number of individual per phylogroup varies considerably over time. One hypothesis that could explain these patterns is the presence of strong marine currents (such as the ACC, Ross Gyre and Weddell Gyre) mixing individuals and thus changing continuously their distribution across the Southern Ocean over time. A similar mechanism has been evoked to explain the current diversity and the distribution of sponges across the Southern Ocean (Downey, Griffiths, Linse, & Janussen, 2012).

Furthermore, within phylogroups, in a few years, the genetic makeup can change significantly. For instance, the genetic composition of the phylogroup C and D changed totally between 1996 and 2008, respectively in the East Weddell Sea and Ross Sea (Fig. 2b, d). Such spatio-temporal variations in term of genetic composition could also be attributed to marine currents that constantly shuffle individuals between populations. Over time, the constant arrival of individuals stemming from different populations continually modify the observed genetic composition within the same geographical location.

Alternatively, a rapid divergence of the genetic pool by genetic drift, or any other population processes, could also explain this rapid genetic turn over. The low effective population sizes highlighted in this study (*Ne* < 10; Table 2) are also consistent with this pattern. Indeed, lower values of *Ne* are associated with increased amount of “apparent” genetic drift. Furthermore, isolated populations are expected to diverge inversely proportional to their effective population sizes. Therefore, when the effective population sizes are extremely low, populations can significantly diverge from each other in just a few generations (Hudson & Coyne, 2002).

The simulations conducted in this study (Fig. 4) show that gene flow from an unsampled reservoir should inflate the *Ne* values estimated with the method of Fu (Fu, 2001). Therefore, due to the extremely low value of *Ne* observed, the hypothesis of a rapid divergence by “apparent” genetic drift is favored here over the hypothesis of a rapid divergence mediated by marine current continually modifying the genetic pool of the populations.

It is also important to bear in mind that our estimates of *Ne* depend directly on the number of generations per year (Fu, 2001). In Table 2, we assumed a generation time of three year, however if for example we assumed a generation time of 30 years, our estimates of *Ne* would increase tenfold. In our case, depending on the assumed value of the generation time, the interpretation of the *Ne* could change accordingly.

Overall, the *Ne* values estimated in this study for the *P. kerguelensis* and *F. mawsoni* are low. However, they are highly abundant in the Southern Ocean (Hemery et al., 2013a; McClintock & Pearse, 1987) which is indicative of a high census size. Such a discrepancy between the census (*Nc*) and effective (*Ne*) population size is commonly observed in nature, especially for marine species with large size (Filatov, 2019). However, the reasons behind it are not always clear (Filatov, 2019; Palstra & Fraser, 2012). At least four biological phenomena could cause *Nc* and *Ne* to be different in *P. kerguelensis* complex (with *Ne* ≪ *Nc*). First, an unequal sex ratio, i.e. an uneven number of males and females contributing to the next generation, lead the rarer sex to have a greater effect on genetic drift which decreases *Ne* compare to *Nc*. Although this hypothesis is plausible, to this day the sex ratio of *P. kerguelensis* and *F. mawsoni* remains unknown. Furthermore, the sex-ratio distortion would need to be extreme to reach such a low *Ne*. Second, a fluctuation of *Nc* over time can affect the ratio between *Ne* and *Nc*. Indeed, an estimate of *Ne* over several generations is the harmonic mean of *Nc* over time. Therefore, if *Nc* changes substantially over time, *Ne* is typically smaller than *Nc*. Fig. 2 indicates that the proportion of *P. kerguelensis* and *F. mawsoni* lineages can vary radically from year to year and thus the fluctuation of *Nc* over time for each lineage can change consequently leading *Ne* to be smaller than *Nc*. Third, the reproductive strategy of *P. kerguelensis* and *F. mawsoni* can participate to reduce *Ne* compared to *Nc*. Indeed, when individuals contribute unequally to the progeny of the next generation, a great proportion of the next generation comes from a small number of individuals which reduces *Ne*. Due to its mode of reproduction, *P. kerguelensis* is particularly incline to have a very reduced number of individuals contributing disproportionately to the progeny. Reproduction occurs synchronously around the months of November and December where males and females spawn and due to the dispersion of gametes by currents, a very small proportion of individuals have their ovocytes fertilized (McClintock & Pearse, 1987). Therefore, only a very small part of the produced larvae contributes effectively to the gene pool. This phenomenon can drastically reduce *Ne* compare to *Nc* and has been reported in different spawning species (Lind, Evans, Knauer, Taylor, & Jerry, 2009). This mode of reproduction can also explain the spatio-temporal genetic turn over mentioned previously. Lastly, due to its haploid and uni-parental nature, the *Ne* of the mtDNA is known to be four times smaller than that of nuclear DNA. Moreover, this effect can be exacerbated by the action of natural selection. Indeed, natural selection can reduce *Ne* and the mtDNA is known to be under positive and negative selection (Meiklejohn, Montooth, & Rand, 2007). Positive natural selection, for example, is expected to reduce the genetic diversity leading to a low *Ne* (Gossmann, Woolfit, & Eyre-Walker, 2011). Indeed, recurrent selective sweeps, related to positive mutations, will reduce the genetic diversity by hitchhiking resulting in a lower *Ne* compared to *Nc*. In this regard, Piganeau & Eyre-Walker (2009) found a strong negative correlation between *Ne* and the strength of the natural selection operating on the mtDNA on non-synonymous mutations. In addition, the non-recombining nature of the mtDNA is expected to make this effect even stronger. There are accumulating evidences suggesting that positive natural selection is acting on echinoderm mitogenomes (Castellana, Vicario, & Saccone, 2011). For example, the strong bias of codon usage observed in the mtDNA of the crinoid species *Florometra serratissima* compared to many other echinoderms, is also in favor of the idea that some sort of natural selection is occurring, though the mechanism behind it is still not fully understood (Scouras & Smith, 2001).

### Tempo of speciation: Promachocrinus kerguelensis challenges the biodiversity pump hypothesis

The mutation rate estimated in this study is approximatively equal to 10^−5^ mutations/site/year and is based on population genetics data. Previous estimate of the mutation rate based also on the COI applied to closely related species (Lessios et al., 1999) used a phylogeny based approach and yielded a mutation rate of roughly 3 order of magnitude lower (≈ 10^−8^ mutations/site/year). Such a discrepancy between mutation rates derived from population data and phylogeny has been already documented (Burridge, Craw, Fletcher, & Waters, 2008; Madrigal et al., 2012) and is called “time dependency of molecular rates” (Ho et al., 2011; Ho, Phillips, Cooper, & Drummond, 2005). For example, in human, the mitochondrial D-loop mutation rate derived from pedigree approaches produced an average value of 8.10^−7^ mutations/site/year whereas phylogenetic estimates yielded value of 2.10^−8^ mutations/site/year (Santos et al., 2005). Likewise, in fishes, Burridge (Burridge et al., 2008) observed a mutation rate of ≈10^−7^ using a population based approach and ≈10^−8^ mutations/site/year from a phylogenetic estimates. The potential causes of the “time scale dependency” are reviewed in Ho *et al.* (2011), but one of the main reason behind it is that population genetics based calibrations lead to estimates reflecting the spontaneous mutation rates, whereas phylogeny based calibrations reflect the substitution rates (i.e. fixed mutations). Therefore, because purifying selection remove the vast majority of the spontaneous deleterious mutations, which constitute a large proportion of them, the spontaneous mutation rate is higher than the substitution rate. Consequently, the rates of molecular evolution decrease with the age of the calibration used to estimate them and it is invalid to assume that a unique molecular clock applies over all time scales (Ho et al., 2005; 2011). A major implication of time dependency is that estimates of recent population divergence times will require re-estimation (Burridge et al., 2008). Therefore, many studies suggesting population isolation related to the Pleistocene glaciations might need to convert these time estimates to more recent ones (i.e. last glacial maximum or Holocene).

Furthermore, the large variation of the mutation rates across the mtDNA regions and between different lineages (Nabholz, Glemin, & Galtier, 2009), leads the mtDNA to strongly deviate from the molecular clock assumption, which is generally assumed in phylogenetic reconstruction. Therefore, estimates of mtDNA mutation rates relying on phylogenetic approach should be taken with caution (Nabholz et al., 2009; Nabholz, Glemin, & Galtier, 2008). Conversely, the mutation rate estimated in this study are quite robust as they are model-free ((Fu, 2001); See also Material and Methods). In the case of *P. kerguelensis* and *F. mawsoni*, the different mutation rates estimated have important consequences on the age of the separation of the different lineages (Fig. 4). Indeed, the estimate from Lessios *et al.* (1999) suggest that the split between the different lineages would have occurred in the last million year (Fig. 4), which is congruent with the hypothesis of the biodiversity pump suggesting that most of the speciation events would have occurred during the Quaternary cycles (Clarke et al., 1992; Clarke & Crame, 1989). Conversely, with our estimate the separation would have occurred in the last 1000 years, implying an extremely recent and rapid diversification for *P. kerguelensis* and *F. mawsoni*.

Explosive radiations have been reported for several taxa (Mahler, Ingram, Revell, & Losos, 2013; Moore & Robertson, 2014; Muschick, Indermaur, & Salzburger, 2012) and are, in general, associated with the Pleistocene glacial cycles (Hawlitschek et al., 2012). In contrast, using the molecular rate estimated in this study, in our case the diversification would have happened during the Holocene. Such a remarkably rapid diversification has been rarely recorded in nature (but see Muschick et al., 2012; Peccoud, Simon, McLaughlin, & Moran, 2009). Furthermore, most of rapid radiations reported so far are adaptive and associated to morphological novelties that allow taxa to exploit separate niches. Taxa involved in rapid diversification are generally ecologically/morphologically highly differentiated (Losos & Miles, 2002). Here, most of the lineages display no obvious morphological differences (except for *Florometra*) or niche differentiation. As a consequence, the radiation within the *P. kerguelensis* complex is probably mainly nonadaptive (Rundell & Price, 2009) or adaptive but related to characters that are challenging to distinguish/observe (physiology or behavior). In this context, it is worth noting that structural genomic changes could also drive taxa to rapid adaptive (Kirkpatrick & Barton, 2006) and nonadaptive (Rowe, Aplin, Baverstock, & Moritz, 2011; Rundell & Price, 2009) radiation and may represent the mechanism underlying the diversification of the *P. kerguelensis* complex.

Finally, the low effective population sizes estimated (Table 2) are congruent with the rapid diversification rate observed. Indeed, Hudson and Coyne (2002) showed that for mtDNA, under the isolation model, two lineages would take between 2 and 3*Ne* generations to become reciprocally monophyletic by lineage sorting. Therefore, using the low effective population size estimated in this study (Table 2), it would take a few hundreds of years for two isolated taxa to reach complete reciprocal monophyly. This result is concordant with the divergence observed in this study and contributes to invalidate the Antarctic “biodiversity pump” for the *Promachocrinus* complex.

## Acknowledgments

The following persons deserve our special thanks for having collected, curated and made available specimens from all around Antarctica: Ty Hibberd (AAD, Hobart), Owen Anderson, David Bowden, Sadie Mills, Kareen Schnabel and Peter Smith (NIWA, Wellington), Stefano Schiaparelli (University of Genoa), Jens Bohn and Eva Lodde (ZSM, Munich), David Barnes, Katrin Linse and Chester Sands (BAS, Cambridge). Our thanks also go to crew and scientists on board various cruises: CEAMARC (IPY project 53), POKERII, TAN08 and AMLR2009 cruises. Funding parties also include three Actions Transversales du MNHN: “Biodiversité actuelle et fossile; crises, stress, restaurations et panchronisme: le message systématique”, “Taxonomie moléculaire: DNA Barcode et gestion durable des collections” and “Biominéralisation”; the French Polar Institute IPEV (travel grants to LGH and ME on REVOLTA); This work was supported by the Consortium National de Recherche en Génomique, and the Service de Systématique Moléculaire (SSM) at the MNHN (USM 2700). Part of the molecular work was also supported by collaboration between the Census of Antarctic Marine Life, the Marine Barcode of Life (MarBOL) project and the Canadian Centre for DNA Barcoding (CCDB). DS was supported by funding of the Alfred P. Sloan Foundation to MarBOL. Laboratory analyses on sequences generated at the CCDB were funded by the Government of Canada through Genome Canada and the Ontario Genomics Institute (2008-OGI-ICI-03). We also gratefully thank Lucile Perrier, Charlotte Tarin, Jose Ignacio Carvajal III Patterson (students) and Céline Bonillo (SSM) for their invaluable help in the molecular lab. This work was also funded by the University of Groningen through a PhD fellowship allocated to Yacine Ben Chehida. We also thank Michael C. Fontaine for the logistical support provided during the PhD of Yacine Ben Chehida.

