## Supplemental Table S1 for "The rapid divergence of the Antarctic crinoid species *Promachocrinus kerguelensis*"

**Table S1. Count of the number of sequences per phylogroup per year and per geographical location**

| Georgia | 1996 | 2000 | 2002 | 2004 | 2005 | 2006 | 2008 | 2009 | 2010 | 2011 |
|---|---|---|---|---|---|---|---|---|---|---|
| A | . | . | . | 1 | . | . | . | . | . | . |
| B | . | . | . | . | . | . | . | . | . | . |
| C | . | . | . | . | . | . | . | . | . | . |
| D | . | . | . | . | . | . | . | . | . | . |
| E | . | . | . | . | . | . | . | . | . | . |
| F | . | . | . | . | . | . | . | . | . | . |
| G | . | . | . | . | . | . | . | . | . | . |

| Sandwich | 1996 | 2000 | 2002 | 2004 | 2005 | 2006 | 2008 | 2009 | 2010 | 2011 |
|---|---|---|---|---|---|---|---|---|---|---|
| A | . | . | . | 9 | . | . | . | . | . | . |
| B | . | . | . | . | . | . | . | . | . | . |
| C | . | . | . | 3 | . | . | . | . | . | . |
| D | . | . | . | . | . | . | . | . | . | . |
| E | . | . | . | . | . | . | . | . | . | . |
| F | . | . | . | . | . | . | . | . | . | . |
| G | . | . | . | . | . | . | . | . | . | . |

| WP | 1996 | 2000 | 2002 | 2004 | 2005 | 2006 | 2008 | 2009 | 2010 | 2011 |
|---|---|---|---|---|---|---|---|---|---|---|
| A | . | . | . | . | . | . | . | . | . | . |
| B | . | . | . | . | . | . | . | . | . | . |
| C | . | . | . | . | . | . | . | . | 12 | . |
| D | . | . | . | . | . | . | . | . | 10 | . |
| E | . | . | . | . | . | . | . | . | 2 | . |
| F | . | . | . | . | . | . | . | . | 2 | . |
| G | . | . | . | . | . | . | . | 10 | 2 | . |

| AS | 1996 | 2000 | 2002 | 2004 | 2005 | 2006 | 2008 | 2009 | 2010 | 2011 |
|---|---|---|---|---|---|---|---|---|---|---|
| A | . | . | . | . | . | . | . | . | . | . |
| B | . | . | . | . | . | . | . | . | . | . |
| C | . | . | . | . | . | . | 3 | . | . | . |
| D | . | . | . | . | . | . | 15 | . | . | . |
| E | . | . | . | . | . | . | . | . | . | . |
| F | . | . | . | . | . | . | 1 | . | . | . |
| G | . | . | . | . | . | . | 3 | . | . | . |

| RS | 1996 | 2000 | 2002 | 2004 | 2005 | 2006 | 2008 | 2009 | 2010 | 2011 |
|---|---|---|---|---|---|---|---|---|---|---|
| A | . | . | . | 85 | . | . | 1 | . | . | . |
| B | . | . | . | 3 | . | . | 37 | . | . | . |
| C | . | . | . | 4 | . | . | 6 | . | . | . |
| D | . | . | . | 27 | . | . | 26 | . | . | . |
| E | . | . | . | 2 | . | . | 74 | . | . | . |
| F | . | . | . | . | . | . | 4 | . | . | . |
| G | . | . | . | 2 | . | . | 26 | . | . | . |

| TAP | 1996 | 2000 | 2002 | 2004 | 2005 | 2006 | 2008 | 2009 | 2010 | 2011 |
|---|---|---|---|---|---|---|---|---|---|---|
| A | . | . | . | . | . | . | 32 | . | . | . |
| B | . | . | . | . | . | . | 54 | . | . | . |
| C | . | . | . | . | . | . | 179 | . | . | . |

| | | | | | | | | | | |
|---|---|---|---|---|---|---|---|---|---|---|
| D | . | . | . | . | . | . | 78 | . | . | . |
| E | . | . | . | . | . | . | 62 | . | . | . |
| F | . | . | . | . | . | . | 9 | . | . | . |
| G | . | . | . | . | . | . | 240 | . | . | . |

| **DS** | 1996 | 2000 | 2002 | 2004 | 2005 | 2006 | 2008 | 2009 | 2010 | 2011 |
|---|---|---|---|---|---|---|---|---|---|---|
| A | . | . | . | . | . | . | . | . | . | . |
| B | . | . | . | . | . | . | . | . | . | . |
| C | . | . | . | . | . | . | . | . | 54 | . |
| D | . | . | . | . | . | . | . | . | 38 | . |
| E | . | . | . | . | . | . | . | . | 22 | . |
| F | . | . | . | . | . | . | . | . | 67 | . |
| G | . | . | . | . | . | . | . | . | 91 | . |

| **KPP** | 1996 | 2000 | 2002 | 2004 | 2005 | 2006 | 2008 | 2009 | 2010 | 2011 |
|---|---|---|---|---|---|---|---|---|---|---|
| A | . | . | . | . | . | . | . | . | . | 120 |
| B | . | . | . | . | . | . | . | . | . | . |
| C | . | . | . | . | . | . | . | . | . | . |
| D | . | . | . | . | . | . | . | . | . | . |
| E | . | . | . | . | . | . | . | . | . | . |
| F | . | . | . | . | . | . | . | . | . | . |
| G | . | . | . | . | . | . | . | . | 28 | . |

| **EWS** | 1996 | 2000 | 2002 | 2004 | 2005 | 2006 | 2008 | 2009 | 2010 | 2011 |
|---|---|---|---|---|---|---|---|---|---|---|
| A | . | . | . | 3 | . | . | . | . | . | . |
| B | 2 | . | . | 1 | . | . | . | . | . | . |
| C | 39 | 37 | . | 24 | . | . | . | . | . | . |
| D | 9 | 2 | . | 2 | 8 | . | . | . | . | . |
| E | 3 | 10 | . | 11 | . | . | . | . | . | . |
| F | . | 4 | . | . | 3 | . | . | . | . | . |
| G | 6 | 3 | . | 14 | 44 | . | 1 | . | . | . |

| **SAW** | 1996 | 2000 | 2002 | 2004 | 2005 | 2006 | 2008 | 2009 | 2010 | 2011 |
|---|---|---|---|---|---|---|---|---|---|---|
| A | . | . | 4 | . | . | . | . | . | . | . |
| B | . | . | . | . | . | . | . | . | . | . |
| C | . | . | 7 | . | . | 3 | . | 32 | . | . |
| D | . | . | . | . | . | 2 | . | 11 | . | . |
| E | . | . | . | . | . | . | . | 25 | . | . |
| F | . | . | . | . | . | 2 | . | . | . | . |
| G | . | 2 | . | . | . | . | . | . | . | . |

| **SAE** | 1996 | 2000 | 2002 | 2004 | 2005 | 2006 | 2008 | 2009 | 2010 | 2011 |
|---|---|---|---|---|---|---|---|---|---|---|
| A | . | . | 2 | . | . | . | . | . | . | . |
| B | . | . | . | . | . | . | . | . | . | . |
| C | . | . | 23 | . | . | 6 | . | . | . | . |
| D | . | . | . | . | . | . | . | . | . | . |
| E | . | . | . | . | . | . | . | . | . | . |
| F | . | . | . | . | . | 1 | . | . | . | . |
| G | . | . | 7 | . | . | . | . | . | . | . |

| Antarctica | 1996 | 2000 | 2002 | 2004 | 2005 | 2006 | 2008 | 2009 | 2010 | 2011 |
|---|---|---|---|---|---|---|---|---|---|---|
| A | . | . | 6 | 98 | . | . | 33 | . | . | 120 |
| B | 2 | . | . | 4 | . | . | 91 | . | . | . |
| C | 39 | 37 | 30 | 31 | . | 9 | 188 | 32 | 66 | . |
| D | 9 | 2 | . | 29 | 8 | 2 | 119 | 11 | 48 | . |
| E | 3 | 10 | . | 13 | . | . | 136 | 25 | 24 | . |
| F | . | 4 | . | . | 3 | 3 | 14 | . | 69 | . |
| G | 6 | 5 | 7 | 16 | 44 | . | 270 | 10 | 121 | . |
